# Functional and structural characterization of allosteric activation of Phospholipase Cε by Rap1A

**DOI:** 10.1101/2020.07.10.191643

**Authors:** Monita Sieng, Arielle F. Selvia, Elisabeth E. Garland-Kuntz, Jesse B. Hopkins, Isaac J. Fisher, Andrea T. Marti, Angeline M. Lyon

**Author notes:** Biotium, 46117 Landing Parkway, Fremont, CA 94538. Corresponding author: Angeline M. Lyon.

## Abstract

Phospholipase Cε (PLCε) is activated downstream of G protein-coupled receptors (GPCRs) and receptor tyrosine kinases (RTKs) through direct interactions with small GTPases, including Rap1A and Ras. While Ras has been reported to allosterically activate the lipase, it is not known whether Rap1A has the same ability, or what its molecular mechanism might be. Rap1A activates PLCε in response to the stimulation of β-adrenergic receptors (β-ARs), translocating the complex to the perinuclear membrane. Because the C-terminal Ras association (RA2) domain of PLCε was proposed to the primary binding site for Rap1A, we first confirmed using purified proteins that the RA2 domain is indeed essential for activation by Rap1A. However, we also showed that the PLCε pleckstrin homology (PH) domain and first two EF hands (EF1/2) are required for Rap1A activation, and identified hydrophobic residues on the surface of the RA2 domain that are also necessary for activation by the GTPase. Finally, small angle X-ray scattering (SAXS) showed that Rap1A binding induces and stabilizes discrete conformational states in PLCε variants that can be activated by the GTPase. This data, together with the recent structure of a catalytically active fragment of PLCε, provide the first evidence that Rap1A, and by extension Ras, allosterically activate the lipase by promoting and stabilizing interactions between the RA2 domain and the PLC core.

## INTRODUCTION

Phospholipase C (PLC) enzymes hydrolyze phosphatidylinositol lipids from cellular membranes in response to diverse cellular signals (1,2). These proteins hydrolyze phosphatidylinositol-4,5-bisphosphate (PIP_2_) at the plasma membrane, producing the second messengers inositol-1,4,5-triphosphate (IP_3_) and diacylglycerol (DAG), which increase Ca^2+^ in the cytoplasm and activate protein kinase C (PKC), respectively. However, some PLC subfamilies, such as PLCε, also hydrolyze other phosphatidylinositol phosphate (PIP) species at internal membranes (3-6). Thus, PLC enzymes regulate multiple pathways from different subcellular locations (1,2).

PLCε is required for maximum Ca^2+^-induced Ca^2+^ release in the cardiovascular system (7,8). In pathological conditions such as heart failure, PLCε expression and activity are increased, promoting overexpression of genes involved in cardiac hypertrophy through a PKC-dependent mechanism (4-6,9,10). Like other PLCs, PLCε contains conserved core domains, including a pleckstrin homology (PH) domain, four EF hand repeats (EF1-4), the catalytic TIM barrel domain, and a C2 domain (1). However, unique N- and C-terminal regulatory regions flank the core. The N-terminal region contains a CDC25 domain that is a guanine nucleotide exchange factor (GEF) for the Rap1A GTPase (11-13), whereas the C-terminal region contains two Ras association (RA) domains, RA1 and RA2 (Figure 1A). Recent functional analysis of a catalytically active PLCε fragment containing the EF3-RA1 domains confirmed that the CDC25, PH, and RA2 domains, as well as EF hands 1/2 (EF1/2) are dispensable for expression and activity. The structure of this fragment revealed that the RA1 domain and the linker connecting the C2 and RA1 domains (C2-RA1) form extensive interactions with EF3/4, the TIM barrel, and C2 domain, suggesting assimilation of the RA1 domain into the catalytic core of this PLC subfamily enzyme (14,15).

**Figure 1.**
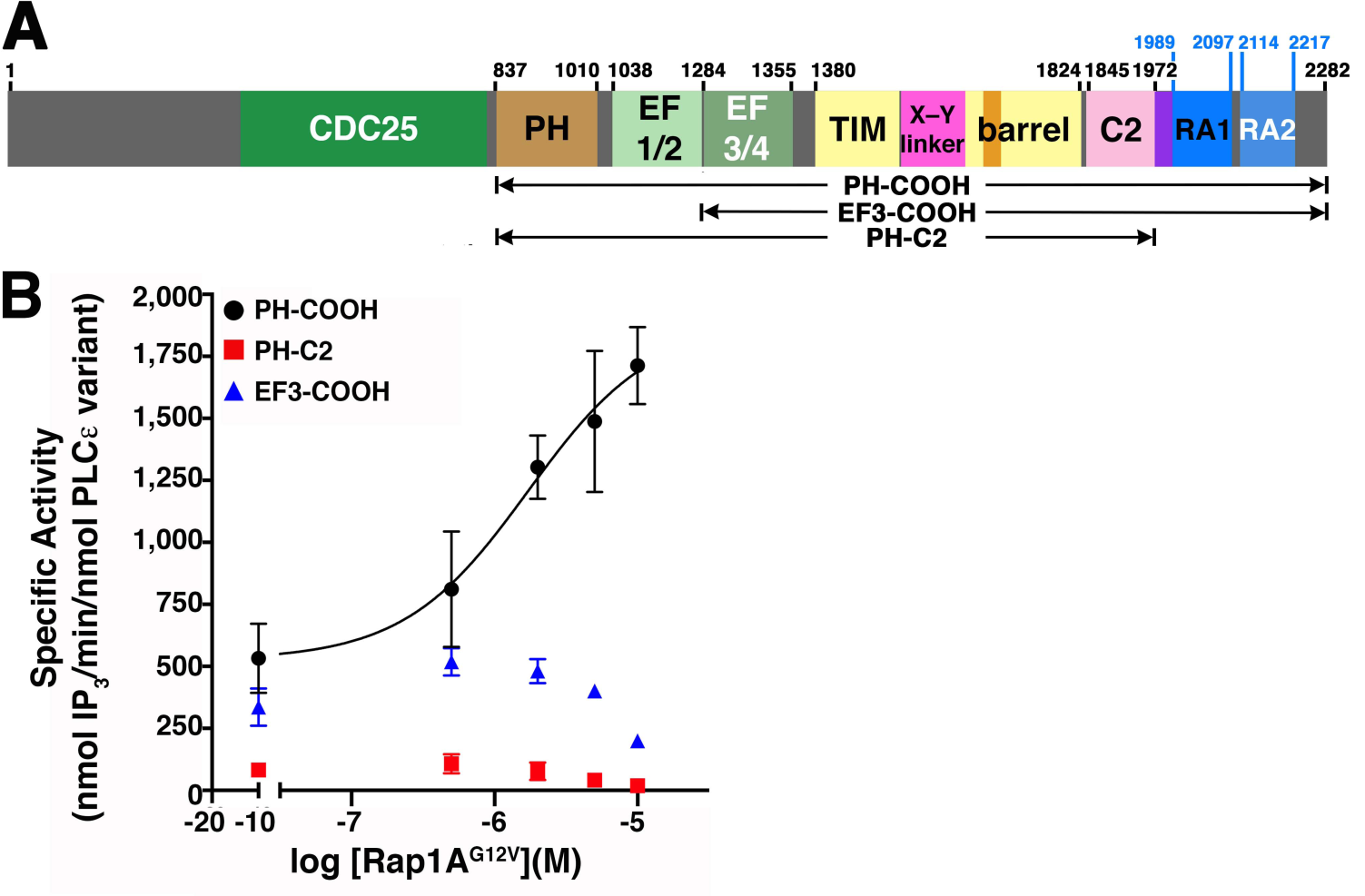
Multiple domains in PLCε are required for Rap1A^G12V^-dependent activation. (A) Domain diagram of rat PLCε. The Y-box insertion and C2-RA1 linker are shown in orange and purple, respectively. Numbers above the diagram correspond to the domain boundaries most relevant to this work, with the variants under study shown below. (B) PLCε PH-COOH (black circles) is activated by Rap1A^G12V^ in a concentration-dependent manner. In contrast, PH-C2 (red squares) and EF3-COOH (blue triangles) are not activated at all concentrations of Rap1A^G12V^ tested. Data represents at least two independent experiments performed in duplicate. Error bars represent SD.

The PLCε RA1 and RA2 domains also contribute to subcellular localization and activation via direct interactions with the scaffolding protein mAKAP, and the Rap1A and Ras GTPases, respectively (1,6,16-18). Of these, the activation of PLCε by Rap1A has been most studied. Stimulation of β-adrenergic receptors in the cardiovascular system activates adenylyl cyclase, increasing cyclic AMP (cAMP), which in turn activates exchange protein activated by cAMP (Epac). Epac catalyzes nucleotide exchange on Rap1A, which binds the RA2 domain, thereby recruiting and allosterically activating PLCε at the Golgi and perinuclear membranes for phosphatidylinositol-4-phosphate (PI4P) hydrolysis (6-10). The GEF activity of PLCε for Rap1A results the formation of a local pool of activated Rap1A, establishing a feed-forward activation loop (19-21). Sustained signaling through this pathway is thought to be one of the key processes underlying pathologic cardiac hypertrophy (3-5,9,10).

Ras-dependent activation of PLCε has been partially characterized, and its mechanism requires both a membrane association and allosteric component (14,17). Although Rap1A is anticipated to share a similar mechanism of activation, it has not yet been shown to allosterically activate PLCε, and the mechanism by which this would occur is not known. We hypothesized that Rap1A binding works in concert with the membrane surface to promote interdomain contacts in PLCε that stabilize a more catalytically competent state, as has been reported for Gα_q_-dependent activation of the related PLCβ enzyme (22). We found that multiple domains of PLCε, in addition to RA2, are required for activation by constitutively active Rap1A and that hydrophobic residues on the surface of the RA2 domain are also essential for Rap1A activation. Finally, we used small angle X-ray (SAXS) scattering to show that Rap1A binding induces and stabilizes large scale structural changes in PLCε that are consistent with the 3D architecture of the enzyme. Together, these results provide new insights into the structure and molecular mechanism of allosteric activation of PLCε by Rap1A.

## RESULTS

### Rap1A-dependent activation of PLCε requires multiple domains of the lipase

Rap1A-dependent activation of PLCε has been demonstrated in cell-based assays, but not using purified components (20,21,23). As full-length PLCε has not been purified in sufficient quantities for biochemical analysis, we relied on the PLCε PH-COOH variant for these studies (Figure 1A)(24), which retains both RA domains and is thus expected to be responsive to Rap1A. In a liposome-based activity assay, the addition of constitutively active and prenylated Rap1A^G12V^ increased the specific activity of PH-COOH ∼3-fold over basal, with a maximum specific activity of 1,900 ± 300 nmol IP_3_/min/nmol PLCε variant (Figure 1B, Table 1, Table S1), which is consistent with the fold activation reported in cell-based assays using full-length PLCε (17,23,25).

**Table 1.** Stability and Rap1A-Dependent Activation of PLCε Variants.

| PLCε variant | T<sub>m</sub> ° C (n) | Basal specific activity<br>(nmol IP<sub>3</sub>/min/nmol<br>PLCε variant) (n)<sup>#</sup> | Fold<br>Activation by<br>Rap1A<sup>G12V</sup> (n) |
| --- | --- | --- | --- |
| PH-COOH* | 51.3 ± 0.72 (6) | 360 ± 120 (9) | 3.0 (4) |
| EF3-COOH | 51.7 ± 0.12 (4) | 450 ± 50 (3) | N/A (3) |
| PH-C2* | 48.3 ± 1.00 (4)<sup>a</sup> | 80 ± 20 (5)<sup>a</sup> | N/A (2) |
| PH-COOH K2150A | 50.5 ± 0.86 (3) | 160 ± 30 (4)<sup>b</sup> | 1.8 (2) |
| PH-COOH K2152A | 49.6 ± 0.79 (3) | 240 ± 20 (3) | 0.9 (3) |
| PH-COOH Y2155A | 49.8 ± 1.06 (3) | 250 ± 40 (5) | 1.4 (3) |
| PH-COOH L2158A | 51.2 ± 0.39 (3) | 270 ± 100 (3) | 1.0 (3) |
| PH-COOH L2192A | 51.5 ± 0.71 (3) | 270 ± 50 (3) | 1.3 (3) |
| PH-COOH F2198A | 51.5 ± 0.45 (3) | 280 ± 100 (3) | 1.2 (3) |

\* T<sub>m</sub> and specific activity for these variants were previously reported, and are included here for comparison (24).
<sup>#</sup>Maximum specific activities were measured at a single time point.
Results are based on one-way ANOVA followed by Dunnett's multiple comparisons test vs. PLCε PH-COOH. Error bars represent SD.
<sup>a</sup> \*\*\*\*p ≤ 0.0002, <sup>b</sup> \*\* p ≤ 0.0028

The RA2 domain is expected to be the primary binding site for Rap1A^G12V^, but other regions of PLCε may also be required for activation. Given that the PLC core domains EF3-C2 are essential for basal activity(26-29), we focused on the roles of the N-terminal PH domain and EF hands 1/2 (EF1/2) in Rap1A-dependent activation. The PLCε EF3-COOH fragment lacks these elements, and has similar stability and basal activity to PH-COOH (Table 1, Figure S1). As a negative control, we also investigated the PH-C2 variant, which has reduced stability and activity relative to PH-COOH (Table 1, Figure S1)(24) and lacks both RA domains. Indeed, PLCε PH-C2 was not activated by Rap1A^G12V^ at any concentration tested, consistent with the absence of the RA2 domain (Figure 1B, Table 1). Surprisingly, PLCε EF3-COOH also showed no activation, indicating that the PLCε PH domain and/or EF1/2 are also necessary for Rap1A-dependent activation (Figure 1B, Table 1, Table S1).

### Hydrophobic residues on the RA2 domain surface are required for Rap1A-dependent activation

Because our data was consistent with multiple PLCε domains contributing to Rap1A^G12V^-dependent activation, we hypothesized that Rap1A allosterically activates PLCε by promoting or stabilizing potentially long range intra- and interdomain interactions within the lipase. As both Ras and Rap1A bind to and activate the enzyme via the RA2 domain, the RA2 domain seems most likely to mediate these interactions. To date, only two residues on the RA2 domain have been characterized with respect activation by GTPases. Mutation of K2150 and/or K2152 (*R. norvegicus* numbering) decrease basal activity and eliminate G protein-dependent activation in cell-based studies (17,23). Based on the structure of activated H-Ras bound to the isolated RA2 domain, K2150 makes an electrostatic interaction with switch II in the GTPase, whereas K2152 is thought to contribute to the local electrostatic environment (14).

We hypothesized that residues involved in allosteric activation on the RA2 domain need to be surface exposed, and would likely be conserved and hydrophobic residues. Using the H-Ras–RA2 structure to model the Rap1A–RA2 interaction (Figure 2A, PDB ID 2C5L (14)), we identified four conserved hydrophobic residues on the surface of the RA2 domain involved in lattice contacts in the crystal structure that did not interact with the GTPase. These residues, Y2155, L2158, L2192, and F2198 (*R. norvegicus* numbering), were individually mutated to alanine in the background of the PLCε PH-COOH variant, and their melting temperatures (T_m_s) and basal activities were determined. As controls, we also expressed, purified, and characterized the PH-COOH K2150 and K2152 point mutants, as they should be insensitive to Rap1A-dependent activation (17,23).

**Figure 2.**
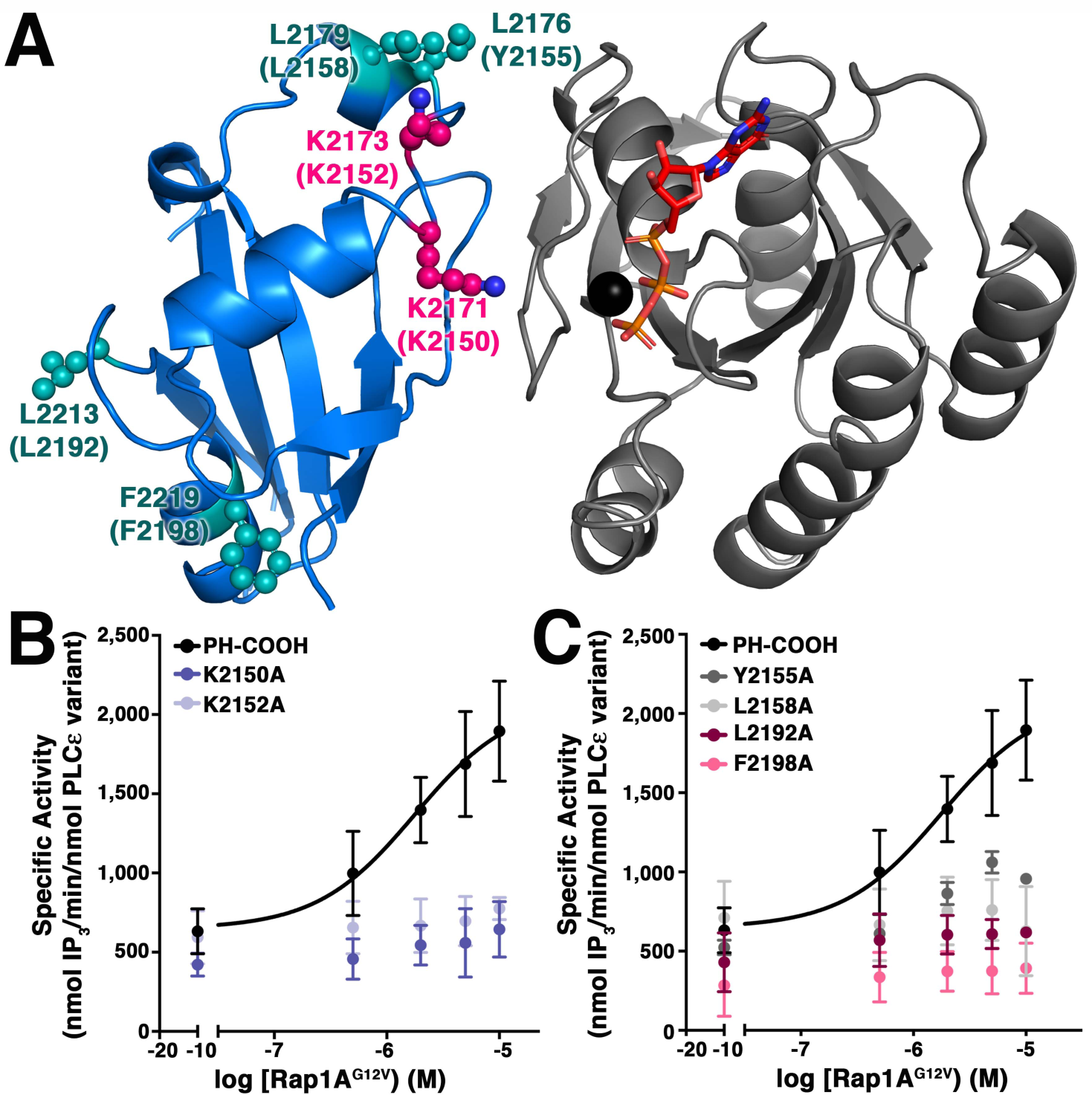
Hydrophobic residues on the surface of the RA2 domain are critical for activation. (A) The structure of H-Ras (gray) bound to the RA2 domain (marine, PDB ID 2C5L(14)) reveals conserved, hydrophobic residues (teal spheres) involved in crystal lattice contacts. K2171 and K2173 (hot pink spheres) were previously reported to be required for Rap1A-dependent activation. *R. norvegicus* residues are in parentheses. GTP is shown in orange sticks, and Mg^2+^ as a black sphere. (B) Mutation of the K2150 or K2152 to alanine eliminates activation by Rap1A^G12V^ *in vitro*, as does (C) mutation of the conserved hydrophobic residues distant from the Rap1A binding surface.

The PH-COOH Y2155A, L2158A, L2192A, and F2198A mutants all had T_m_ values comparable to that of PH-COOH (15,30), and basal specific activities within ∼2-fold of PH-COOH (Table 1, Figure S1) demonstrating they are properly folded. K2150A and K2152A also had T_m_ values comparable to that of PH-COOH, but K2150A had ∼2-fold lower basal activity, consistent with previous reports (Table 1, Figure S1)(17,23). We then tested the ability of Rap1A^G12V^ to activate the point mutants in a liposome-based activity assay. The K2150 and K2152A mutants were insensitive to activation by Rap1A^G12V^, consistent with their proposed role in binding GTPases (Figure 2B, Table 1) (17,23). Mutation of the hydrophobic surface residues also decreased or eliminated Rap1A^G12V^-dependent activation at all concentrations tested (Figure 2C, Table 1, Table S1). Thus, these hydrophobic residues appear to play a critical role in this mechanism.

### Rap1A^G12V^ binding to PLCε induces and stabilizes unique conformational states

Our domain deletion and site-directed mutagenesis analyses identified roles for the PH domain, EF1/2, and the hydrophobic residues on the RA2 surface in Rap1A^G12V^-dependent activation. To gain structural insight into how these elements contribute to activation by Rap1A^G12V^, we used SAXS to compare the solution structures of the PLCε PH-COOH and EF3-COOH variants alone and in complex with Rap1A^G12V^. These variants were chosen because they formed stable complexes with Rap1A^G12V^ that could be isolated by size exclusion chromatography, but only PH-COOH has increased lipase activity upon binding of the GTPase (Figure 1, Table 1).

We first compared the SAXS solution structure of the Rap1A^G12V^–PH-COOH complex with that of PLCε PH-COOH. We previously showed this variant is a globular protein with extended features likely due to the flexibly connected PH, EF1/2, and RA2 domains (Figures 3, 4A, Table 2, Table S2, Figure S2)(24,31). The Rap1A^G12V^–PH-COOH sample was complicated, and contained three minor components that partially overlapped the major complex peak in the SEC-SAXS elution profile (Figure S2). To identify the region corresponding to Rap1A^G12V^–PH-COOH, evolving factor analysis (EFA) was used to deconvolute the data (Figure S3)(32,33). The Rap1A^G12V^–PH-COOH complex had an R_g_ of 42.4 ± 0.12 Å (Figure 3D, E) and a D_max_ of ∼165 Å, with a largely globular structure with some extended features (Figure 3). However, further analysis of the samples, shown in the dimensionless Kratky plot, revealed substantial conformational changes due to Rap1A^G12V^ binding (Figure 4B). In this plot, compact, globular proteins have bell-shaped curves that converge to the qR_g_ axis at high values, whereas elongated and more rigid structures exhibit curves that extend out to higher qR_g_, and highly flexible structures do not converge at all (34). The data for the Rap1A^G12V^–PH-COOH complex shows the overall structure is more compact and/or less flexible than PH-COOH alone, as evidenced by the curve being more bell-shaped and converging to zero at lower values of qRg (Figure 4B).

**Table 2.** SAXS parameters of PLCε PH-COOH and EF3-COOH in complex with Rap1A^G12V^.

|  | **PH-COOH<sup>1</sup>** | **Rap1A<sup>G12V</sup>–**<br>**PH-COOH** | **EF3-COOH** | **Rap1A<sup>G12V</sup>–**<br>**EF3-COOH** |
| --- | --- | --- | --- | --- |
| *Guinier analysis* |  |  |  |  |
| $I(0)$ (Arb.) | $64.92 \pm 0.21$ | $1.66 \pm 0.003$ | $78.10 \pm 0.43$ | $26.96 \pm 0.50$ |
| $R_g$ (Å) | $42.7 \pm 0.17$ | $42.4 \pm 0.12$ | $41.9 \pm 0.43$ | $36.8 \pm 0.50$ |
| $q$ min (Å<sup>-1</sup>) | 0.0174 | 0.0043 | 0.0084 | 0.0094 |
| $q$ range (Å<sup>-1</sup>) | 0.0174 - 0.0314 | 0.0043 - 0.0306 | 0.0084 - 0.0309 | 0.0094 - 0.352 |
| *P(r) analysis* |  |  |  |  |
| $I(0)$ (Arb.) | $65.96 \pm 0.21$ | $1.67 \pm 0.03$ | $79.43 \pm 0.63$ | $27.81 \pm 0.21$ |
| $R_g$ (Å) | $44.79 \pm 0.19$ | $43.70 \pm 0.16$ | $44.48 \pm 0.61$ | $39.47 \pm 0.43$ |
| $D_{max}$ (Å) | 162 | 165 | 175 | 145 |
| Porod volume (Å<sup>-3</sup>) | 191,000 | 237,000 | 179,000 | 178,000 |
| $q$ range (Å<sup>-1</sup>) | 0.0174 - 0.308 | 0.0042 - 0.350 | 0.0087 – 0.387 | 0.0052 – 0.366 |

<sup>1</sup>The data for the PLCε PH-COOH variant was previously published, and is included for comparison (24).

**Figure 3.**
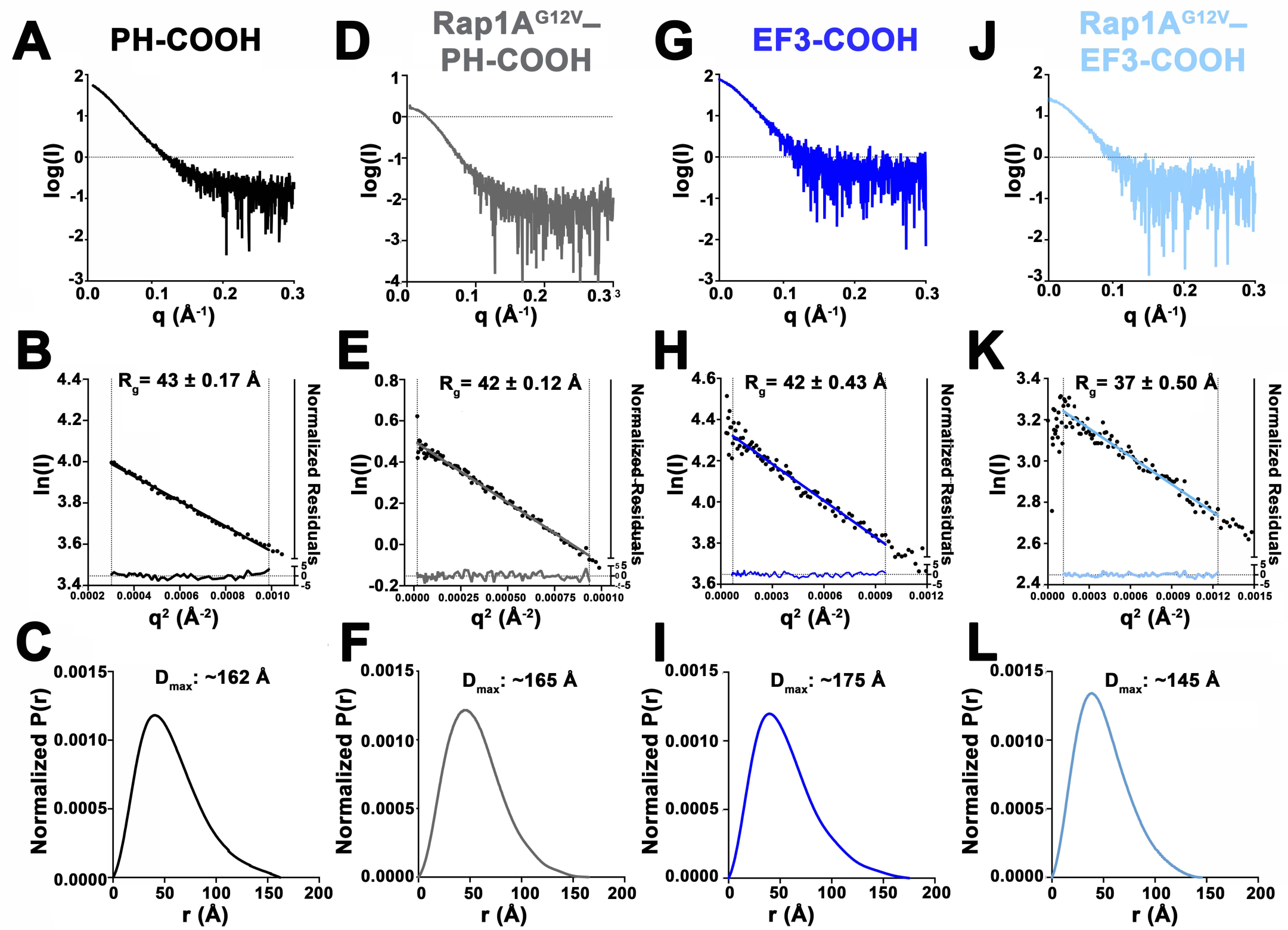
Rap1A^G12V^ binding to PLCε PH-COOH or EF3-COOH stabilizes different conformational states. The (A) scattering profile for PLCε PH-COOH and (B) Guinier plot demonstrate the variant is monomeric and monodisperse in solution. The (C) pair-distance distribution function is consistent with a largely globular protein with some extended features. The (D) scattering profile and (E) Guinier plot for the Rap1A^G12V^–PH-COOH complex is also consistent with a monodisperse complex. The (F) pair-distance distribution function shows a more compact structure upon the binding of Rap1A^G12V^. PLCε EF3-COOH is similar to PH-COOH in solution, as evidenced by its (G) scattering profile, (H) Guinier plot, and (I) pair-distance distribution function. The Rap1A^G12V^–EF3-COOH complex does not have elevated lipase activity but is still monodisperse in solution as shown in (J) the scattering profile and (K) Guinier plot. However, the (L) shape of the pair-distance distribution function reveals the complex is more globular than EF3-COOH alone, and more compact, as evidenced by the smaller D_max_. The data for the PLCε PH-COOH variant is included for comparison (24).

**Figure 4.**
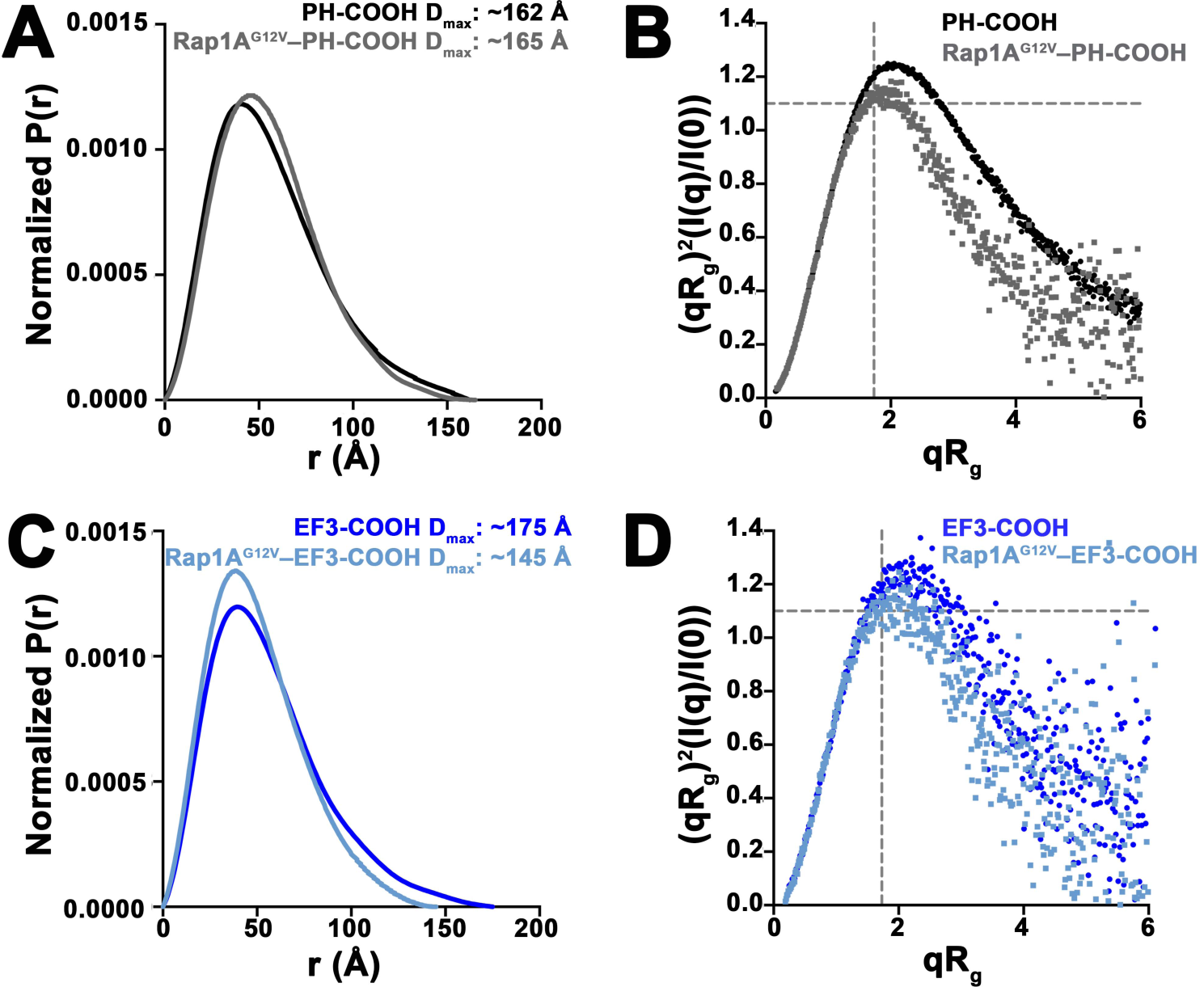
Normalized pair-distance and dimensionless Kratky plots for PLCε variants alone and in complex with Rap1A^G12V^. (A) The normalized P(r) functions for PH-COOH and Rap1A^G12V^–PH-COOH are similar with D_max_ values of ∼162 Å and ∼165 Å, respectively. (B) Comparison of PH-COOH (black circles) and Rap1A^G12V^–PH-COOH (gray squares) show the complex is more compact and globular than PH-COOH alone, as evidenced by the more bell-shaped curve. (C) The normalized P(r) functions for EF3-COOH and Rap1A^G12V^–EF3-COOH reveal binding of Rap1A^G12V^ induces substantial conformational changes that lead to a more compact structure. This is further supported by the ∼30 Å decrease in D_max_ for the Rap1A^G12V^–EF3-COOH complex. (D) Comparison of EF3-COOH (blue circles) and Rap1A^G12V^–EF3-COOH (light blue squares). Rap1A^G12V^ binding induces conformational changes that result in a more compact and globular solution structure.

We next compared the solution structures of PLCε EF3-COOH alone and in complex with Rap1A^G12V^. EF3-COOH was also monomeric and monodisperse in solution, with an R_g_ of 41.9 ± 0.43 Å and a D_max_ of ∼175 Å (Figure 3G-I, Table 2, Table S2, Figure S2). Its P(r) function is also consistent with the protein having a mostly globular structure with a modest degree of extendedness and/or flexibility, consistent with the flexibly connected RA2 domain (Figures 3I, 4C). Surprisingly, the solution structure of the Rap1A^G12V^–EF3-COOH complex differed substantially from EF3-COOH. The complex had an R_g_ of 36.8 ± 0.5 Å (Figure 3J, K, Table S2, Figure S2), and its P(r) function revealed a much more compact and globular structure, as shown in the more bell-shaped curve and most apparently in the ∼30 Å decrease in D_max_ to ∼145 Å (Figures 3, 4, Table 2). This is further highlighted in the dimensionless Kratky plot, which clearly show that Rap1A^G12V^ binding induces conformational changes that result in a more compact, stable structure (Figure 4D). Because EF3-COOH is not activated by Rap1A^G12V^ (Figure 1, Table 1), this more condensed state may correspond to a nonproductive conformation of the complex, or a state in which it is incompetent for activation, due to loss of the PH domain and EF1/2.

## DISCUSSION

The PLCε RA domains are highly similar in structure, but have different functional roles in the enzyme (1,6,15-18). The RA1 domain, together with the C2-RA1 linker, forms extensive contacts with EF hands 3/4 (EF3/4), the TIM barrel, and the C2 domain that are important for stability and activity (15). RA1 also interacts with muscle-specific A-kinase anchoring protein (mAKAP), a scaffolding protein at the Golgi, helping localize PLCε to internal membranes (18). The contribution of the RA2 domain to basal activity is unclear, as its deletion has been reported to either activate, inhibit, or have no minimal impact on basal activity (14,15,17). The RA2 domain is the primary binding site for activated Rap1A and Ras GTPases, as deletion of the domain or mutation of two highly conserved lysines (*R. norvegicus* PLCε K2150 and K2152, Figure 2A) eliminates G protein-stimulated activation in cells (14,17,19). Interestingly, mutation of K2150 alone decreases basal activity ∼50% in cells (17,19). NMR and biochemical studies have shown RA2 is flexibly connected to RA1, and does not stably associate with the PLCε core (14,15). However, how GTPase binding to this domain is translated into increased lipase activity is poorly understood. Given that all known activators PLCε are lipidated, membrane localization is certainly one aspect of the activation mechanism. However, membrane association alone is insufficient to fully stimulate lipase activity. For example, a PLCε variant bearing a -CAAX motif at its C-terminus for constitutive plasma membrane localization had increased lipase activity, but this was stimulated an additional ∼4-fold in the presence of activated Ras, suggesting that the activation mechanism mediated by small GTPases must also have an allosteric component (14).

In this work, we used a series of purified PLCε domain deletion variants and point mutants to investigate the allosteric component of the Rap1A-dependent activation mechanism. We have shown that constitutively activate Rap1A binds to and increases the lipase activity of PLCε PH-COOH *in vitro* to a similar extent as full-length PLCε in cells (Figure 1)(20,21,23). We also found the PH domain and EF1/2 are required for Rap1A-dependent activation (Figure 1, Table 1). These findings support a model in which the binding of Rap1A is a collaborative event involving multiple domains of PLCε.

We next sought to identify conserved residues in the RA2 domain, which is of known structure, that could be involved in intramolecular interactions with the PLCε core upon Rap1A binding. Guided by the previously determined crystal structure of the H-Ras–RA2 complex (Figure 2A, PDB ID 2C5L(14)), we generated a model of the Rap1A–RA2 interaction and identified four conserved, solvent-exposed, hydrophobic resides on RA2 that are distant from the predicted Rap1A binding site. Mutation of Y2155, L2158, L2192, or F2198 to alanine eliminated Rap1A^G12V^-dependent activation (Figure 2, Table 1). One explanation for these results is that the conserved, hydrophobic residues on the RA2 surface interact with or stabilize intramolecular contacts between the RA2 domain and the PLCε core. These intramolecular interactions may function to communicate the fact that a GTPase is bound to the rest of the lipase.

Our study identifies roles for the PLCε PH domain, EF1/2, and conserved hydrophobic residues on the RA2 surface as being critical for Rap1A-dependent activation. These domains are distant in the primary structure of PLCε (Figure 1A), but are situated in relatively close spatial proximity based on the recent structure of the PLCε EF3-RA1 fragment (15). To gain structural insights into how these structural elements could contribute to activation, we used SAXS to compare the solution structures of PLCε PH-COOH and EF3-COOH alone and in complex with Rap1A^G12V^. This comparison allows identification of large-scale conformational changes, changes in shape (globular vs. extended), and differences in flexibility (Figures 4, 5, Table 2, Table S2). The PLCε variants alone had similar globular structures with some extended/flexible features, consistent with the presence of at least one flexibly connected domain. The binding of Rap1A^G12V^ to either variant induced conformational changes that resulted in more ordered and less flexible structures (Figures 3, 4). However, these two Rap1A-bound complexes differ substantially from one another, as Rap1A^G12V^ binding to EF3-COOH decreased the maximum diameter of the complex by ∼20 Å and stabilized a more compact, globular structure (Figures 3, 4, Table 2). Because the lipase activity of EF3-COOH was not increased by Rap1A, this solution structure cannot represent the active state. In contrast, Rap1A^G12V^ binding to PH-COOH did not appreciably change the maximum diameter of the complex, but did lead to a structure that is more globular and less flexible than PH-COOH alone (Figures 3, 4). These results indicate that Rap1A binding may stabilize the interactions between the Rap1A-bound RA2 domain and the PLCε core.

**Figure 5.**
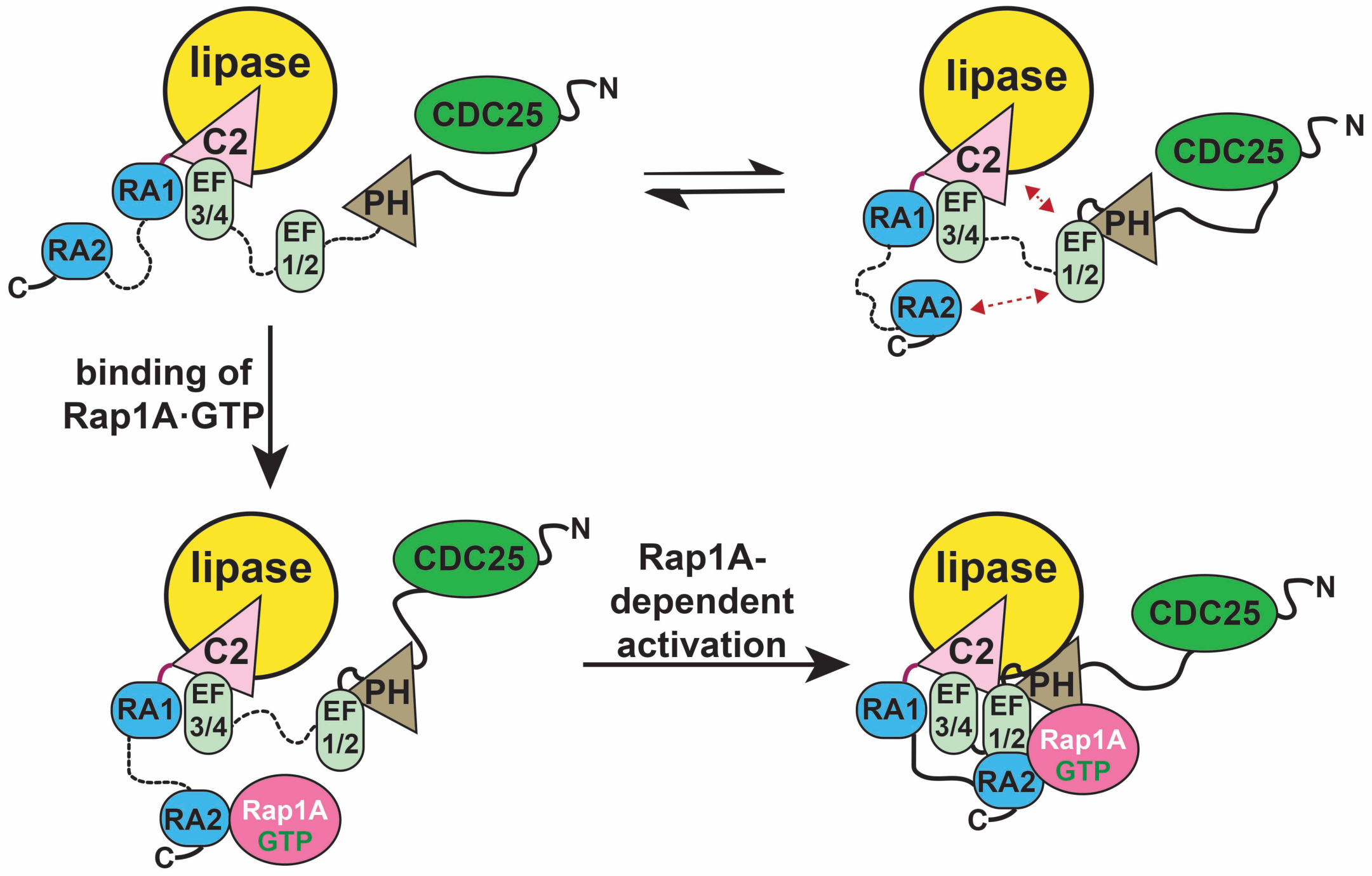
A model for activation of PLCε by Rap1A. (Top) PLCε exists in multiple conformational states in solution. The PH domain, EF1/2, and RA2 domains are flexibly connected to the rest of the enzyme, as indicated by the dashed black lines, and interact transiently with one another under basal conditions (24). (Bottom left) Activated Rap1A binds to its high affinity binding site on the PLCε RA2 domain. However, this interaction is insufficient on its own to activate lipase activity. (Bottom right) The Rap1A–RA2 complex also interacts with a site on the PLCε core, potentially formed by the PH domain and EF hands, which mediate Rap1A-dependent activation.

Overall, our results provide the first direct structural evidence about the nature of the allosteric component of Rap1A-dependent (and potentially Ras-dependent) activation of PLCε, and reveal that activation involves substantial conformational changes within the lipase. Whereas the PLCε RA2 domain is required for activation, this process also appears to be dependent on the PH domain, EF1/2, and hydrophobic surface residues on the RA2 domain. These observations are consistent with the following model (Figure 5). Because the RA2 domain is flexibly tethered to RA1, it may transiently interact with rest the of PLCε core in solution. These interactions are insufficient to alter the thermal stability or increase basal activity. When Rap1A binds to the RA2 domain, this complex could stabilize interactions with other domains in PLCε. This most likely occurs via the hydrophobic residues present on the surface of RA2 with the PH domain and/or EF1/2, based on the location of the N- and C-termini in the crystal structure of PLCε (15). Finally, the membrane itself contributes to Rap1A-dependent activation in several possible ways, such as through a membrane-induced allosteric event, and/or by stabilizing a unique and fully activated Rap1A–PLCε complex. Although the regions responsible for membrane association in PLCε have not yet been identified, the enzyme is able to at least transiently interact with the membrane, given its measurable basal activity (17). Finally, Rap1A is prenylated, and thus, along with any other membrane binding element, likely helps to orient the lipase active site at the membrane for maximum lipid hydrolysis. Future studies that provide higher resolution insights into the interactions between the RA2 domain and the PLCε core, alone and in complex with activated Rap1A, will be essential steps in elucidating a complete picture of this process and new opportunities for therapeutic development that target activation of PLCε by small GTPases.

## EXPERIMENTAL PROCEDURES

### Protein expression, purification, and mutagenesis of PLCε variants

cDNAs encoding N-terminally His-tagged *R. norvegicus* PLCε variants were subcloned into pFastBac HTA (PH-COOH: residues 837-2281, PH-C2: 832-1972, and EF3-COOH: 1284-2281). Site-directed mutagenesis in the PH-COOH background was performed using the QuikChange Site-Directed Mutagenesis Kit (Stratagene) or the Q5 Site-Directed Mutagenesis Kit (NEB). All subcloned PLCε variants contained an N-terminal His-tag and TEV cleavage site, and were sequenced over the entire coding region. The proteins were expressed and purified as previously described (24), with some modifications for the PLCε EF3-COOH used in the SAXS experiments. Briefly, after elution from an Ni-NTA column, PLCε EF3-COOH was incubated with 5% w/w TEV protease to remove the N-terminal His-tag and dialyzed overnight against 1.5 L of buffer containing 20 mM HEPES pH 8.0, 50 mM NaCl, 2 mM DTT, 0.1 mM EDTA, and 0.1 mM EGTA at 4 °C. The dialysate was applied to Roche cOmplete NiNTA resin or a GE HisTrap, and the flow-through containing the TEV-cleaved EF3-COOH was collected and passed over the column two more times. The protein in the collected flow-through was then purified as previously described (24).

### Expression and purification of prenylated Rap1A^G12V^·GTP

cDNA encoding N-terminally His-tagged constitutively active *H. sapiens* Rap1A (Rap1A^G12V^) was subcloned into pFastBac HTA. The protein were expressed in baculovirus-infected High5 cells. Cell pellets were resuspended in lysis buffer containing 20 mM HEPES pH 8.0, 100 mM NaCl, 10 mM β-mercaptoethanol, 0.1 mM EDTA, 10 mM NaF, 20 mM AlCl_3_, 0.1 mM LL, 0.1 mM PMSF, and 20 μM GTP and lysed via dounce on ice. The lysate was centrifuged for 1 h at 100,000 x *g*, and the pellet was resuspended in lysis buffer supplemented with 1% sodium cholate, dounced on ice, and solubilized at 4 °C for 1 h. The sample was then centrifuged for 1 h at 100,000 x *g*, and the supernatant was diluted two-fold with lysis buffer.

His-tagged Rap1A^G12V^ was loaded on a Ni-NTA column pre-equilibrated with lysis buffer, and first washed with lysis buffer containing 10 mM imidazole and 0.2% cholate, followed by a second wash with lysis buffer supplemented with 10 mM imidazole and 10 mM CHAPS (3-[(3-cholamidopropyl)dimethylammonio]-1-propanesulfonate). The protein was eluted with lysis buffer containing 250 mM imidazole pH 8.0 and 10 mM CHAPS. The column was washed with lysis buffer containing 0.2% cholate, followed by lysis buffer supplemented with 10 mM CHAPS. His-tagged Rap1A^G12V^ was then concentrated and applied to tandem Superdex S200 columns pre-equilibrated with G protein S200 buffer (20 mM HEPES pH 8.0, 50 mM NaCl, 1 mM MgCl_2_, 2 mM DTT, 10 mM CHAPS, and 20 μM GTP). Fractions containing purified protein were identified by SDS-PAGE, pooled, concentrated, and flash frozen in liquid nitrogen. Rap1A^G12V^ used for activation assays was purified in modified G protein S200 buffer containing 1 mM CHAPS.

For SAXS experiments, His-tagged Rap1A^G12V^ was incubated with 5% w/w TEV protease and dialyzed overnight in 1.5 L of dialysis buffer containing 20 mM HEPES pH 8.0, 50 mM NaCl, 1 mM MgCl_2_, 10 mM β-mercaptoethanol, 1 mM CHAPS, and 20 μM GTP at 4 °C. The dialysate was applied to a Ni-NTA column, and the flow-through containing cleaved Rap1A^G12V^ was collected and passed over the column two more times. The flow-through was collected, concentrated to 1 mL, applied to tandem Superdex S200 columns and purified as described above.

### Differential scanning fluorimetry (DSF)

Melting temperatures (T_m_) of PLCε variants were determined as previously described (24,30). A final concentration of 0.5 mg/mL was used for each PLCε variant. At least three independent experiments were performed in duplicate.

### PLCε activity assays

All activity assays were carried out using [^3^H]-PIP_2_ as the substrate. Basal activity of PLCε variants was measured as previously described (24). Briefly, 200 μM phosphatidylethanolamine, 50 μM PIP_2_, and ∼4,000 cpm [^3^H]-PIP_2_ were mixed, dried under nitrogen, and resuspended by sonication in buffer containing 50 mM HEPES pH 7, 80 mM KCl, 2 mM EGTA, and 1 mM DTT. Enzyme activity was measured at 30 °C in 50 mM HEPES pH 7, 80 mM KCl, 15 mM NaCl, 0.83 mM MgCl_2_, 3 mM DTT, 1 mg/mL bovine serum albumin (BSA), 2.5 mM EGTA, 0.2 mM EDTA, and ∼500 nM free Ca^2+^. PLCε PH-COOH and EF3-COOH were assayed at a final concentration of 0.075 ng/μL and PH-C2 at 0.1-1ng/μL (24). The PH-COOH K2150A, K2152A, Y2155A, L2158A, L2192A, and F2198A mutants were assayed at a final concentration of 0.5 ng/μL. Control reactions contained everything except free Ca^2+^. Reactions were quenched by addition of 200 μL 10 mg/mL BSA and 10% (w/v) ice-cold trichloroacetic acid and centrifuged. Free [^3^H]-IP_3_ in the supernatant was quantified by scintillation counting. All assays were performed at least three times in duplicate.

Rap1A^G12V^-dependent increases in PLCε lipase activity were measured using the same approach with some modifications. The liposomes were first incubated with increasing concentrations of Rap1A^G12V^·GTP in 50 mM HEPES pH 7.0, 3 mM EGTA, 1 mM EDTA, 100 mM NaCl, 5 mM MgCl_2_, 3 mM DTT, and 390 μM CHAPS at 30 °C for 30 min. The reaction was initiated by addition of the PLCε variant, incubated at 30 °C for 8 min, and processed as described above. All activation assays were performed in duplicate with protein from at least two independent purifications.

### Formation and isolation of the Rap1A^G12V^–PLCε variant complexes

A 1:3 or 1:5 molar ratio of PLCε PH-COOH or EF3-COOH to Rap1A^G12V^, supplemented with 0.5 mM CaCl_2_, was incubated on ice for 30 min before being applied to a Superdex S200 column pre-equilibrated with complex S200 buffer (20 mM HEPES pH 8, 50 mM NaCl, 2 mM DTT, 0.1 mM EDTA, 0.1 mM EGTA, 1 mM MgCl_2_, 0.5 mM CaCl_2_ and 40 μM GTP). Fractions containing the purified complex were identified by SDS-PAGE, pooled, and concentrated for use in SAXS experiments.

### SAXS Data Collection and Analysis

PLCε PH-COOH was previously characterized by SAXS (24). PLCε EF3-COOH, the Rap1A^G12V^–PLCε PH-COOH complex, and the Rap1A^G12V^–PLCε EF3-COOH complex were diluted to final concentrations of 2-3 mg/mL in S200 buffer (EF3-COOH) or complex S200 buffer, and centrifuged at 16,000 x *g* for 5 min at 4 °C prior to data collection. Size exclusion chromatography (SEC)-SAXS was performed at the BioCAT beamline at Sector 18 of the Advanced Photon Source (Table S1).

Protein samples were eluted from a Superdex 200 Increase 10/300 GL column using an ÄKTA Pure FPLC (GE Healthcare) at a flow rate of 0.7 mL/min. The eluate passed through a UV monitor followed by a SAXS flow cell consisting of a quartz capillary. The data were collected in two different setups at the beamline. The PH-COOH, EF3-COOH, and EF3-COOH complex data were collected in a 1.5 mm ID quartz capillary with 10 μm walls. The PH-COOH complex data was collected in a 1.0 mm ID quartz capillary with 50 μm using the coflow sample geometry (35) to prevent radiation damage. Scattering intensity was recorded using Pilatus3 × 1M detector (Dectris) placed ∼3.7 m from the sample using 12 KeV X-rays (1.033 Å wavelength) and a beam size of 160 x 75 μm, giving an accessible q range of ∼0.004 Å^-1^ to 0.36 Å^-1^. Data was collected every 2 s with 0.5 s exposure times. Data in regions flanking the elution peak were averaged to create buffer blanks which were subsequently subtracted from exposures selected from the elution peak to create the final scattering profiles (Figure S2). BioXTAS RAW 1.4.0(33) was used for data processing and analysis. For the Rap1A^G12V^–PLCe PH-COOH complex, the four components present in the sample were deconvoluted using evolving factor analysis (EFA, Figure S3)(32)) as implemented in BIOXTAS RAW(33). The radius of gyration (R_g_) of individual frames were plotted with the scattering chromatograms, which plot integrated intensity of individual exposures as a function of frame number, and used to help determine appropriate sample ranges for subtraction. PRIMUS (36)was used to calculate the R_g_, I_(0)_, and D_max_ for both samples. GNOM (37) was used within PRIMUS to generate the pair distance distribution (P(r)) functions via an indirect Fourier transform (IFT) method.

Graphical plots were generated from buffer-subtracted averaged data (scattering profile and Guinier plots)(38) or IFT data (P(r) plots) and plotted using GraphPad Prism v.8.0.1. SAXS data are presented in accordance with the publication guidelines for small angle scattering data(39).

### Statistical Methods

GraphPad Prism v.8.0.1 was used to generate all plots. One-way ANOVA was performed with Prism v.8.0.1 followed by Dunnett’s post-hoc multiple comparisons vs. PLCε PH-COOH, as noted in figure captions. All error bars represent standard deviation.

## SUPPORTING INFORMATION

Supporting Table 1. Minimum and Maximum Specific Activities under Rap1A^G12V^ activating conditions

Supporting Table 2. SAXS Molecular Weight Estimations for PLCε PH-COOH and EF3-COOH alone and in complex with Rap1A^G12V^.

Supporting Figure 1. Characterization of the PLCε domain deletion variants and RA2 point mutants

Supporting Table 3. SAXS Data Collection and Analysis Parameters

Supporting Figure 2. Size exclusion chromatography (SEC-SAXS) scattering chromatograms for PLCε variants alone and in complex with Rap1A^G12V^.

Supporting Figure 3 Deconvolution of the Rap1A^G12V^–PH-COOH elution peak using evolving factor analysis (EFA).

## AUTHOR CONTRIBUTIONS

M.S. and A.M.L. designed the experimental approach. M.S., E.E.G.-K, A.F.S., and A.T.M. cloned, expressed, and purified all PLCε and Rap1A proteins. M.S., E.E.G.-K., and I.J.F. performed DSF and activity assays. M.S. and A.F.S. isolated Rap1A–PLCε complexes for SAXS analysis. J.B.H. assisted with SAXS experiments, data analysis, and processing. M.S., E.E.G.-K., J.B.H., and A.M.L. wrote the manuscript.

## ACKNOWLEDGMENTS

We thank S. Chakravarthy (APS BioCAT) for assistance with SAXS data collection and analysis. Use of the Advanced Photon Source, an Office of Science User Facility operated for the U. S. Department of Energy (DOE) Office of Science by Argonne National Laboratory, was supported by the U.S. DOE under Contract Number DE-AC02-06CH11357. This project was supported by grant 9 P41 GM103622 from the National Institute of General Medical Sciences of the National Institutes of Health. Use of the Pilatus 3 1M detector was provided by grant 1S10OD018090-01 from NIGMS.

The content is solely the responsibility of the authors and does not necessarily represent the official views of the National Institute of General Medical Sciences or the National Institutes of Health.

## ABBREVIATIONS

PLC: Phospholipase C
RA: Ras association
DAG: diacylglycerol
IP_3_: inositol-1,4,5-triphopshate
PKC: protein kinase C
CHAPS: 3-[(3-cholamidopropyl)dimethyl-ammonio]-1-propane sulfonate
GEF: guanine exchange factor
GPCR: G protein-coupled receptor
RTK: receptor tyrosine kinase
cAMP: cyclic AMP
Epac: exchange protein activated by cAMP
PIP: phosphatidylinositol phosphate
PIP_2_: phosphatidylinositol (4,5)-bisphosphate
PE: phosphatidylethanolamine
PH: pleckstrin homology
TIM: triose-phosphate isomerase
RA: Ras association
PI_4_P: phosphatidylinositol-4-phosphate
DSF: differential scanning fluorimetry
SAXS: small angle X-ray scattering
SEC: size exclusion chromatography

## REFERENCES

1. Kadamur, G., and Ross, E. M. (2013) Mammalian phospholipase C. Annu Rev Physiol 75, 127–154

2. Gresset, A., Sondek, J., and Harden, T. K.(2012) The phospholipase C isozymes and their regulation. Subcell Biochem 58, 61–94

3. de Rubio, R. G., Ransom, R. F., Malik, S., Yule, D. I., Anantharam, A., and Smrcka, A. V. (2018) Phosphatidylinositol 4-phosphate is a major source of GPCR-stimulated phosphoinositide production. Science Signaling 11

4. Nash, C. A., Brown, L. M., Malik, S., Cheng, X., and Smrcka, A. V. (2018) Compartmentalized cyclic nucleotides have opposing effects on regulation of hypertrophic phospholipase Cepsilon signaling in cardiac myocytes. J Mol Cell Cardiol

5. Nash, C. A., Wei, W., Irannejad, R., and Smrcka, A. V. (2019) Golgi localized beta1-adrenergic receptors stimulate Golgi PI4P hydrolysis by PLCepsilon to regulate cardiac hypertrophy. eLife 8

6. Smrcka, A. V., Brown, J. H., and Holz, G. G.(2012) Role of phospholipase Cε in physiological phosphoinositide signaling networks. Cell Signal 24, 1333–1343

7. Oestreich, E. A., Malik, S., Goonasekera, S. A., Blaxall, B. C., Kelley, G. G., Dirksen, R. T., and Smrcka, A. V. (2009) Epac and phospholipase Cε regulate Ca^2+^ release in the heart by activation of protein kinase Cε and calcium-calmodulin kinase II. J Biol Chem 284, 1514–1522

8. Oestreich, E. A., Wang, H., Malik, S., Kaproth-Joslin, K. A., Blaxall, B. C., Kelley, G. G., Dirksen, R. T., and Smrcka, A. V.(2007) Epac-mediated activation of phospholipase Cε plays a critical role in beta-adrenergic receptor-dependent enhancement of Ca^2+^ mobilization in cardiac myocytes. J Biol Chem 282, 5488–5495

9. Zhang, L., Malik, S., Pang, J., Wang, H., Park, K. M., Yule, D. I., Blaxall, B. C., and Smrcka, A. V. (2013) Phospholipase cepsilon hydrolyzes perinuclear phosphatidylinositol 4-phosphate to regulate cardiac hypertrophy. Cell 153, 216–227

10. Wang, H., Oestreich, E. A., Maekawa, N., Bullard, T. A., Vikstrom, K. L., Dirksen, R. T., Kelley, G. G., Blaxall, B. C., and Smrcka, A. V. (2005) Phospholipase C ε modulates β-adrenergic receptor-dependent cardiac contraction and inhibits cardiac hypertrophy. Circ Res 97, 1305–1313

11. Jin, T. G., Satoh, T., Liao, Y., Song, C., Gao, X., Kariya, K., Hu, C. D., and Kataoka, T.(2001) Role of the CDC25 homology domain of phospholipase Cε in amplification of Rap1-dependent signaling. J Biol Chem 276, 30301–30307

12. Satoh, T., Edamatsu, H., and Kataoka, T.(2006) Phospholipase Cε guanine nucleotide exchange factor activity and activation of Rap1. Methods Enzymol 407, 281–290

13. Dusaban, S. S., Kunkel, M. T., Smrcka, A. V., and Brown, J. H. (2015) Thrombin Promotes Sustained Signaling and Inflammatory Gene Expression through the CDC25 and Ras Associating Domains of Phospholipase C-epsilon. J Biol Chem 290, 26776–26783

14. Bunney, T. D., Harris, R., Gandarillas, N. L., Josephs, M. B., Roe, S. M., Sorli, S. C., Paterson, H. F., Rodrigues-Lima, F., Esposito, D., Ponting, C. P., Gierschik, P., Pearl, L. H., Driscoll, P. C., and Katan, M. (2006) Structural and mechanistic insights into ras association domains of phospholipase C ε. Mol Cell 21, 495–507

15. Rugema, N. Y., Garland-Kuntz, E. E., Sieng, M., Muralidharan, K., Van Camp, M. M., O’Neill, H., Mbongo, W., Selvia, A. F., Marti, T., Everly, A., McKenzie, E., and Lyon, A. M. (2020) Structure of phospholipase Cε reveals an integrated RA1 domain and previously unidentified regulatory elements. Communications Biology 3, 445

16. Wing, M. R., Bourdon, D. M., and Harden, T. K. (2003) PLC-ε: a shared effector protein in Ras-, Rho-, and Gαβγ-mediated signaling. Mol Interv 3, 273–280

17. Kelley, G. G., Reks, S. E., Ondrako, J. M., and Smrcka, A. V. (2001) Phospholipase Cε: a novel Ras effector. EMBO J 20, 743–754

18. Zhang, L., Malik, S., Kelley, G. G., Kapiloff, M. S., and Smrcka, A. V. (2011) Phospholipase C ε scaffolds to muscle-specific A kinase anchoring protein (mAKAPβ) and integrates multiple hypertrophic stimuli in cardiac myocytes. J Biol Chem 286, 23012–23021

19. Kelley, G. G., Kaproth-Joslin, K. A., Reks, S. E., Smrcka, A. V., and Wojcikiewicz, R. J.(2006) G-protein-coupled receptor agonists activate endogenous phospholipase Cε and phospholipase Cβ3 in a temporally distinct manner. J Biol Chem 281, 2639–2648

20. Edamatsu, H., Satoh, T., and Kataoka, T.(2006) Ras and Rap1 activation of PLCepsilon lipase activity. Methods Enzymol 407, 99–107

21. Song, C., Satoh, T., Edamatsu, H., Wu, D., Tadano, M., Gao, X., and Kataoka, T. (2002) Differential roles of Ras and Rap1 in growth factor-dependent activation of phospholipase C epsilon. Oncogene 21, 8105–8113

22. Lyon, A. M., Dutta, S., Boguth, C. A., Skiniotis, G., and Tesmer, J. J. (2013) Full-length Galpha(q)-phospholipase C-beta3 structure reveals interfaces of the C-terminal coiled-coil domain. Nat Struct Mol Biol 20, 355–362

23. Kelley, G. G., Reks, S. E., and Smrcka, A. V.(2004) Hormonal regulation of phospholipase Cε through distinct and overlapping pathways involving G12 and Ras family G-proteins. Biochem J 378, 129–139

24. Garland-Kuntz, E. E., Vago, F. S., Sieng, M., Van Camp, M., Chakravarthy, S., Blaine, A., Corpstein, C., Jiang, W., and Lyon, A. M.(2018) Direct observation of conformational dynamics of the PH domain in phospholipases C and beta may contribute to subfamily-specific roles in regulation. J Biol Chem 293, 17477–17490

25. Khrenova, M. G., Mironov, V. A., Grigorenko, B. L., and Nemukhin, A. V. (2014) Modeling the role of G12V and G13V Ras mutations in the Ras-GAP-catalyzed hydrolysis reaction of guanosine triphosphate. Biochemistry 53, 7093–7099

26. Essen, L. O., Perisic, O., Cheung, R., Katan, M., and Williams, R. L. (1996) Crystal structure of a mammalian phosphoinositide-specific phospholipase C δ. Nature 380, 595–602

27. Zhang, W., and Neer, E. J. (2001) Reassembly of phospholipase C-β2 from separated domains: analysis of basal and G protein-stimulated activities. J Biol Chem 276, 2503–2508

28. Seifert, J. P., Wing, M. R., Snyder, J. T., Gershburg, S., Sondek, J., and Harden, T. K. (2004) RhoA activates purified phospholipase C-ε by a guanine nucleotide-dependent mechanism. J Biol Chem 279, 47992–47997

29. Seifert, J. P., Zhou, Y., Hicks, S. N., Sondek, J., and Harden, T. K. (2008) Dual activation of phospholipase C-ε by Rho and Ras GTPases. J Biol Chem 283, 29690–29698

30. Mezzasalma, T. M., Kranz, J. K., Chan, W., Struble, G. T., Schalk-Hihi, C., Deckman, I. C., Springer, B. A., and Todd, M. J. (2007) Enhancing recombinant protein quality and yield by protein stability profiling. J Biomol Screen 12, 418–428

31. Korasick, D. A., and Tanner, J. J. (2018) Determination of protein oligomeric structure from small-angle X-ray scattering. Protein science : a publication of the Protein Society 27, 814–824

32. Meisburger, S. P., Taylor, A. B., Khan, C. A., Zhang, S., Fitzpatrick, P. F., and Ando, N.(2016) Domain Movements upon Activation of Phenylalanine Hydroxylase Characterized by Crystallography and Chromatography-Coupled Small-Angle X-ray Scattering. Journal of the American Chemical Society 138, 6506–6516

33. Hopkins, J. B., Gillilan, R. E., and Skou, S.(2017) BioXTAS RAW: improvements to a free open-source program for small-angle X-ray scattering data reduction and analysis. Journal of applied crystallography 50, 1545–1553

34. Durand, D., Vives, C., Cannella, D., Perez, J., Pebay-Peyroula, E., Vachette, P., and Fieschi, F. (2010) NADPH oxidase activator p67(phox) behaves in solution as a multidomain protein with semi-flexible linkers. J Struct Biol 169, 45–53

35. Kirby, N., Cowieson, N., Hawley, A. M., Mudie, S. T., McGillivray, D. J., Kusel, M., Samardzic-Boban, V., and Ryan, T. M. (2016) Improved radiation dose efficiency in solution SAXS using a sheath flow sample environment. Acta crystallographica. Section D, Structural biology 72, 1254–1266

36. Konarev, P. V., Volkov, V. V., Sokolova, A. V., Koch, M. H. J., and Svergun, D. I. (2003) PRIMUS: a Windows PC-based system for small-angle scattering data analysis. Journal of applied crystallography 36, 1277–1282

37. Svergun, D. I. (1992) Determination of the Regularization Parameter in Indirect-Transform Methods Using Perceptual Criteria. Journal of applied crystallography 25, 495–503

38. Franke, D., Petoukhov, M. V., Konarev, P. V., Panjkovich, A., Tuukkanen, A., Mertens, H. D. T., Kikhney, A. G., Hajizadeh, N. R., Franklin, J. M., Jeffries, C. M., and Svergun, D. I. (2017) ATSAS 2.8: a comprehensive data analysis suite for small-angle scattering from macromolecular solutions. Journal of applied crystallography 50, 1212–1225

39. Trewhella, J., Duff, A. P., Durand, D., Gabel, F., Guss, J. M., Hendrickson, W. A., Hura, G. L., Jacques, D. A., Kirby, N. M., Kwan, A. H., Perez, J., Pollack, L., Ryan, T. M., Sali, A., Schneidman-Duhovny, D., Schwede, T., Svergun, D. I., Sugiyama, M., Tainer, J. A., Vachette, P., Westbrook, J., and Whitten, A. E. (2017) 2017 publication guidelines for structural modelling of small-angle scattering data from biomolecules in solution: an update. Acta crystallographica. Section D, Structural biology 73, 710–728

