## Supplementary material for "Functional and structural characterization of allosteric activation of Phospholipase Cε by Rap1A": Rap1A_BioRxiv_suppinfo_v1.pdf (from another version of this preprint: version 1, 2020-07-10)

Present Addresses: [#]Sequlite Genomics, 6773 Sierra Ct, Dublin, CA 94568

*Corresponding author: Angeline M. Lyon

**Supporting Table 1.** Minimum and Maximum Specific Activities under Rap1A$^{G12V}$ Activation Conditions

| PLCε variants | Minimum activity ± SD (nmol $IP_3$/min/nmol PLCε variant) | Maximum Rap1A$^{G12V}$-stimulated activity (nmol $IP_3$/min/nmol PLCε variant) (n) |
|---|---|---|
| PH-COOH | 630 ± 100 | 1,900 ± 300 (4) |
| PH-C2 | 70 ± 30 | 24 ± 9 (3) |
| EF3-COOH | 335 ± 80 | 200 ± 15 (2) |
| PH-COOH K2150A | 420 ± 70 | 540 ± 200 (2) |
| PH-COOH K2152A | 590 ± 200 | 700 ± 100 (3) |
| PH-COOH Y2155A | 520 ± 50 | 640 ± 30 (3) |
| PH-COOH L2158A | 710 ± 200 | 780 ± 70 (3) |
| PH-COOH L2192A | 430 ± 200 | 960 ± 30 (3) |
| PH-COOH F2198A | 380 ± 100 | 630 ± 300 (3) |

These specific activities are determined at a single time point.

**Supporting Table 2.** SAXS Molecular Weight Estimations for PLCε PH-COOH and EF3-COOH alone and in complex with Rap1A$^{G12V}$

| | PH-COOH[1] | Rap1A$^{G12V}$–PH-COOH | EF3-COOH | Rap1A$^{G12V}$–EF3-COOH |
|---|---|---|---|---|
| *P(r) analysis* | | | | |
| Porod Volume MW (kDa), (ratio to expected) | 130 (0.76) | 200 (1.0) | 150 (1.3) | 120 (0.88) |
| Volume of Correlation MW (kDa), (ratio to expected) | 120 (0.71) | 160 (0.82) | 120 (1.1) | 100 (0.73) |
| Calculated MW (kDa) | 165.5 | 189.8 | 112.8 | 134.2 |

[1] The data for the PLCε PH-COOH variant was previously published[22], and is included here for comparison.

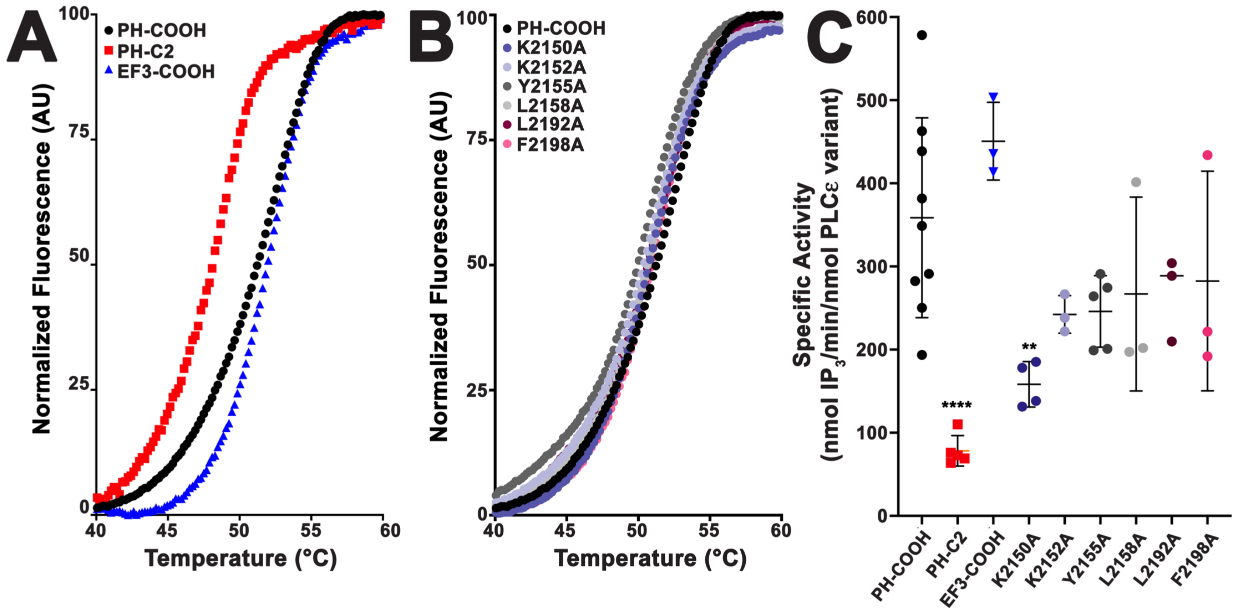


**Supporting Figure 1.** Characterization of the PLCε domain deletion variants and RA2 point mutants. (A) Representative thermal denaturation curves of the PLCε domain deletion variants used in this study. PH-COOH is shown in black circles (23), PH-C2 in red squares (23), and EF3-COOH in blue triangles. (B) Representative thermal denaturation curves the PH-COOH RA2 point mutants. PH-COOH is shown as in (A), K2150A in navy blue circles, K2152 in periwinkle circles, Y2155A in gray circles, L2158A in light gray circles, L2192A in maroon circles, and F2198A in pink circles. (C) Scatter plot of the basal specific activities of the PLCε variants. All assays were performed at least three times in duplicate, and error bars represent SD. Significance was determined using a one-way ANOVA followed by Dunnett's multiple comparisons test vs. PLCe PH-COOH. **** $p \leq 0.0002$, ** $p \leq 0.0028$.

**Supporting Table 3.** SAXS Data Collection and Analysis Parameters

| | |
|---|---|
| *(a)* SAXS data collection parameters | |
| Instrument | BioCAT facility at the Advanced Photon Source beamline 18ID with Pilatus3 X 1M (Dectris) detector |
| Wavelength (Å) | 1.033 |
| Beam size ($mm^2$) | 160 (h) x 75 (v) |
| Camera length (m) | 3.7 |
| $q$-measurement range ($Å^{-1}$) | 0.004-0.4 |
| Absolute scaling method | N/A or Glassy Carbon, NIST SRM 3600 |
| Basis for normalization to constant counts | To incident intensity, by ion chamber counter |
| Method for monitoring radiation damage | Automated frame-by-frame comparison of relevant regions using CORMAP[35] implemented in BioXTAS RAW |
| Exposure time, number of exposures | 0.5 s exposure time with 2 s exposure period (0.5 s on, 1.5 s off) of entire SEC elution |
| Sample configuration | SEC-SAXS. Separation by size using ÄKTA Pure with a Superdex 200 Increase 10/300 GL column. Some SAXS data measured in a 1.5 mm ID quartz capillary while others used a sheath-flow cell[24] with effective path length 0.542 mm. |
| Sample temperature (˚C) | 20 |
| *(b)* Software employed for SAXS data reduction, analysis and interpretation | |
| SAXS data reduction | Radial averaging; BioXTAS RAW 1.4.0[25] and ATSAS[29] used for frame comparison, averaging, and subtraction |
| Basic analysis: Guinier, M.W., P(r) | BioXTAS RAW 1.4.0[25] used for Guinier fit and molecular weight; GNOM[36] used for P(r) function |

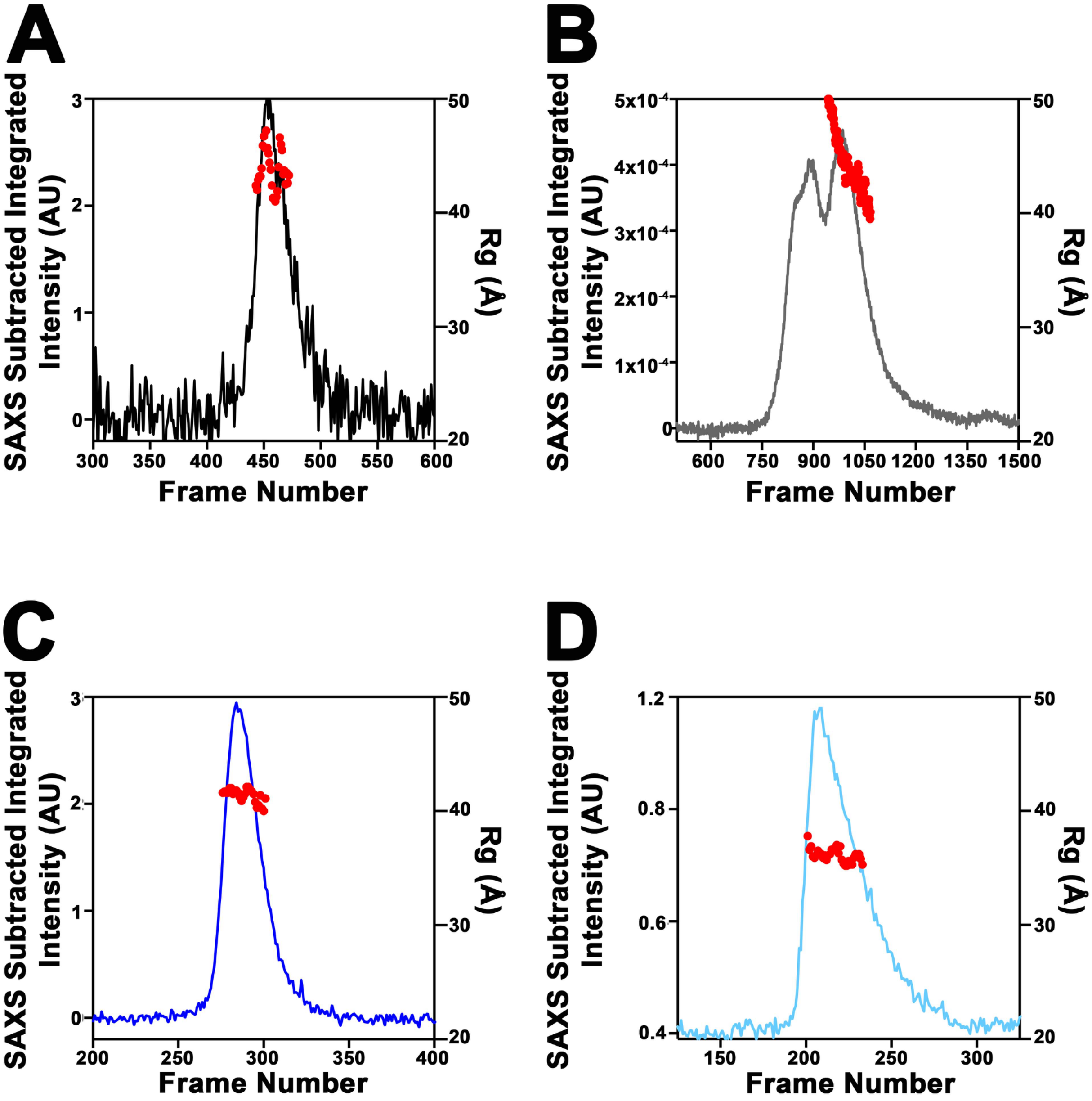


**Supporting Figure 2.** Size exclusion chromatography (SEC-SAXS) scattering chromatograms for PLCε variants alone and in complex with Rap1A$^{G12V}$. Each chromatogram shows the SAXS subtracted integrated intensity and $R_g$ as a function of exposure (frame) number for (A) PLCε PH-COOH (23), (B) Rap1A$^{G12V}$–PH-COOH, (C) PLCε EF3-COOH, and (D) Rap1A$^{G12V}$–EF3-COOH. Red circles correspond to $R_g$.

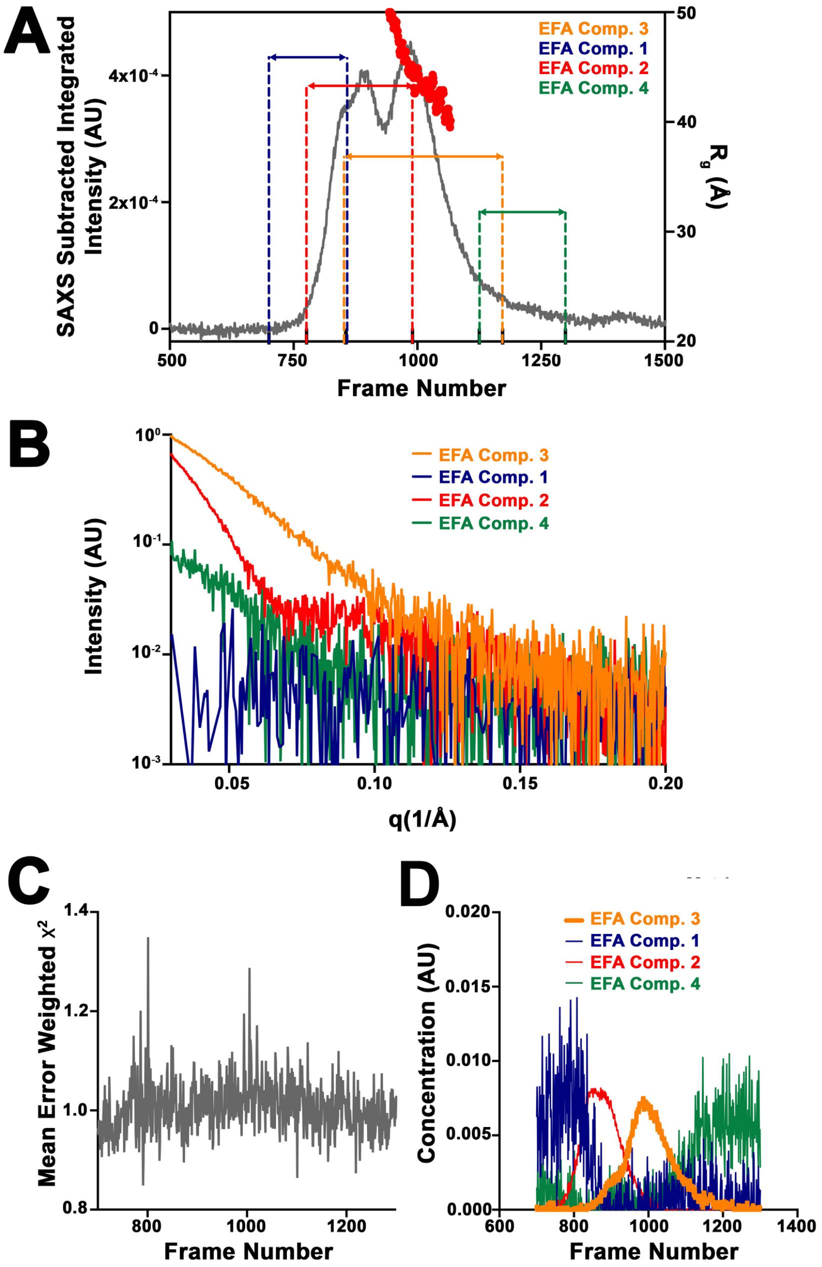


**Supporting Figure 3.** Deconvolution of the Rap1A$^{G12V}$–PH-COOH elution peak using evolving factor analysis (EFA). (A) Size exclusion chromatogram (SEC-SAXS) scattering chromatogram showing the SAXS subtracted integrated intensities and $R_g$ as a function of frame number. Four components were identified in the Rap1A$^{G12V}$–PH-COOH sample (EFA Comp. 1-4), and their corresponding ranges are shown in dashed lines. EFA Comp. 3 (orange) corresponds to Rap1A$^{G12V}$–PH-COOH. (B) Overlaid SAXS scattering curves for each component identified by EFA. (C) Mean error weighted $\chi^2$ of the EFA deconvolution. Values near 1 show a good fit of the deconvolution results to the original data. (D) Area normalized concentration profiles extracted from EFA for each component, colors correspond to region colors in A. This shows clearly where in the elution each component comes from, and that each component represents a relatively pure species.
